# Free-living marine nematodes diversity at Ponta Delgada-São Miguel (Azores archipelago, North-East Atlantic Ocean): first results from shallow soft-bottom habitats

**DOI:** 10.1101/2020.09.09.289918

**Authors:** Alberto de Jesús Navarrete, Víctor Aramayo, Anitha Mary Davidson, Ana Cristina Costa

## Abstract

Contrasting (sand, algae, rocky-dominated, mixed) benthic habitats were sampled to characterize marine nematode diversity inhabiting surface sediments in São Miguel (Azores, North-East Atlantic Ocean) in July 2019. Nematodes were extracted from the surface layer of sediments and morphologically identified using light microscopy. Nematode taxonomy was based on living/fresh specimens) to ensure a suitable recognition of morphological traits. Our results provide a preliminary checklist of free-living marine nematode genera from 21 intertidal and sublittoral sandy beach sites along the coast of São Miguel island, Azores archipelago, Portugal. The nematode fauna was represented by 4 orders, 21 families, and 43 genera. *<u>Cyatholaimus</u>, <u>Desmodora</u>* and *<u>Daptonema</u>* had two morphospecies each. Enoplida was represented by 8 families and 13 genera, while Chromadorida by 7 families and 18 genera, the latter corresponding to the most diverse nematode group. Monhysterida had 5 families and 10 genera and Araeolaimida was represented by a single 1 family and 2 genera. The most common genera (i.e., accounting for 75% of all organisms) included *<u>Adoncholaimus</u>* (most abundant genus, 32 specimens), *<u>Axonolaimus</u>* (18), *<u>Cyatholaimus</u>* (17), *<u>Enoploides</u>* (13), *<u>Rhabdocoma</u>*, and *<u>Acanthopharynx</u>* (11). *<u>Viscosia</u>* and *<u>Enoplolaimus</u>* were represented by 7 specimens, whilst *<u>Halalaimus</u>, <u>Desmoscolex</u>, <u>Monophostia</u>, <u>Daptonema</u>*, and *<u>Theristus</u>* obtained only 6 each. The dominant nematode taxa of São Miguel island have been commonly previously reported in other coastal habitats including sandy beaches. They can be considered typical meiofaunal components of intertidal sandy beaches. Many of the nematode morphotypes found in São Miguel island could represent new species to science. As far as we know, this is the first report on free-living nematodes for São Miguel Island and for marine shallow water in the Azores. Our findings will serve as an import baseline for future research aiming to improve our understating of nematode communities in volcanic islands such as São Miguel in the Azores archipelago.

## Resumo

Diversos habitats bentónicos (areia, algal, rochoso e misto) foram amostrados para caracterizar a diversidade de nematodes marinhos que habitam o sedimento superficial em São Miguel (Açores, Oceano Atlântico Nordeste) em Julho 2019. Os nematodes foram extraídos da camada superficial de sedimento e identificados morfologicamente usando microscopia óptica. A identificação taxonómica realizou-se com base na observação de especímenes vivos e/ou frescos para garantir o reconhecimento adequado dos caracteres morfológicos. Os resultados providenciam uma lista taxonómica preliminar dos nematodes marinhos de vida livre de 21 locais intertidais e subtidais de praias arenosas ao longo da costa da ilha de São Miguel, no arquipélago dos Açores, Portugal. A fauna de nematodes obtida encontrou-se representada por 4 ordens, 21 famílias e 43 géneros. Os géneros *<u>Cyatholaimus</u>, <u>Desmodora</u>* e *<u>Daptonema</u>* apresentaram 2 morfoespécies cada. A ordem Enoplida representou-se por 8 famílias e 13 géneros, Chromadorida por 7 familias e 18 géneros, a última correspondendo ao grupo de nematodes mais diverso. Monhysterida apresentou 5 famílias e 10 géneros e Araeolaimida estava representada por uma única família e 2 géneros. O género mais comum (contribuindo para 75% de todos os organismos amostrados) incluiu *<u>Adoncholaimus</u>* (o género mais abundante, 32 especímenes), *<u>Axonolaimus</u>* (18), *<u>Cyatholaimus</u>* (17), *<u>Enoploides</u>* (13), *<u>Rhabdocoma</u>*, e *<u>Acanthopharynx</u>* (11). *<u>Viscosia</u>* e *<u>Enoplolaimus</u>* foram representados por 7 especímenes, enquanto que *<u>Halalaimus</u>, <u>Desmoscolex</u>, <u>Monophostia</u>, <u>Daptonema</u>*, e *<u>Theristus</u>* foram representados por apenas 6 indivíduos cada. Os *taxa* dominates de nematodes marinhos na ilha de São Miguel têm sido comumente reportados para outros habitats costeiros incluindo praias arenosas. Podem ser considerados um componente típico da meiofauna de praias arenosas intertidais. Muitos dos morfotipos de nematodes marinhos encontrados na ilha de São Miguel podem representar novas espécies para a ciência. Tanto quanto sabemos, este é o primeiro registo de nematodes marinhos para a ilha de São Miguel e a primeira lista de nematos marinhos de baixa profundidade para os Açores. Estes resultados servirão como uma baseline importante para futuras investigações com o objectivo de melhorar a nossa compreensão sobre as comunidades de nematodes em ilhas vulcânicas como a ilha de São Miguel no arquipélago dos Açores.

**Palavras chave: nematodes marinhos de vida livre, meiofauna, habitats bentónicos intertidais, sedimentos vulcânicos.**

## Introduction

Shallow marine environments such as sandy beaches are among the most appreciated ecosystems by humankind not only for their recreational and economic value but also for the contribution of ecosystem services they provide like: sediment storage and transport, wave dissipation and habitat for benthic organisms (Short, 1996; McLachlan & Brown, 2006). The very condition of sandy beaches makes them look like lifeless habitats, in terms of macrofauna, however, these environments are home to a diverse and abundant interstitial fauna composed mainly by meiofauna; animals smaller than 500 µm and larger than 35 µm (McLachlan & Jaramillo 1995; Giere, 2009).

Free-living nematodes are one of the most abundant meiofaunal organisms in sedimentary environments (>80% of total community abundance), where they play a pivotal role in the recycling of organic matter and energy flow, thus acting as a link between the microfauna and the benthic macrofauna as well as higher trophic groups (e.g. fishes; Bhadury *et al*., 2015). Nematodes are considered ubiquitous and are found in almost all environments (Heip *et al*., 1985), where their spatial distribution and abundance levels are often shaped by several environmental factors. However, sediment type plays a fundamental role in nematode distribution. Previous studies have shown that some nematode families such as Desmodoridae, Microlaimidae, Camacolaimidae, and Ironidae are often dominant in sandy substrates, while in finer sediments such as silts or clays, Anoplostomidae, Comesomatidae, and Linhomoeidae are usually the dominant ones (Sharma & Webster, 1983; Gourbault & Warwick, 1994; Nicholas & Hodda 1999; Gheskiere *et al*., 2004; Urban-Malinga *et al*., 2004; Calles *et al*., 2005; Hourston *et al*., 2005; Moreno *et al*., 2006; de Jesús-Navarrete, 2007; Mundo-Ocampo *et al*., 2007; Bhadury *et al*., 2015).

It is recognized that research on marine nematodes at European coastal habitats is well advanced with a better understanding of their spatio-temporal patterns, including taxonomic keys for marine nematodes from that region (Platt & Warwick, 1983; 1988; Warwick *et al*., 1998; Lambshead & Boucher, 2003). Numerous studies focused on nematode communities have been carried out in European sandy beaches (Vincx *et al*., 1990; Nicholas & Hodda, 1999; Gheskiere *et al*., 2004; Urban-Malinga *et al*., 2004; Calles *et al*., 2005; Semprucci *et al*., 2013). Unfortunately, our understanding of nematode species distribution in tropical regions is still scarce (Bhadury *et al*., 2015). Some papers have been published above marine nematodes of Canary Islands, (Riera *et al*. 2005; 2014), however, we have very little knowledge of nematode communities from remote volcanic islands such as the Azores archipelago. Nematode studies in this area have solely focused on deep-sea hydrothermal and seamount habitats (Zeppilli *et al*., 2013; 2015; Bellec *et al*., 2018), whereas no reports are available for nematode assemblages in shallow sandy habitats. Overall, nematodes have been a neglected group in local marine diversity studies, mainly due to lack of local taxonomic expertise. (see Costello *et al*., 2006).

In order to contribute to the current knowledge of the diversity at the Azores Archipelago as well as to the diversity of sandy environments in volcanic islands, the present study aims to identify the free-living nematofauna inhabiting surface sediments from shallow soft-bottom habitats at São Miguel island.

## Materials and Methods

### Study area

Like the other islands in the Azores archipelago, São Miguel is a volcanic-originated island, characterized by unique flora and fauna (Borges *et al*., 2020). Being the largest island in this region (Borges *et al*., 2020) exhibits several shallow benthic habitats (e.g. open sandy beaches, intertidal pools, rocky shores). Although some inventorial efforts have been oriented to terrestrial and marine (large) fauna (e.g. mammals, seabirds), less is known about the local marine invertebrate fauna (in particular, the tiny, meiofauna-sized component).

Samples analyzed in the present study were collected during the International Workshop VW-Summer School held at the University of the Azores on July 15-24, 2019. This workshop was focused on the diversity of local meiofaunal groups.

### Sampling strategy

Sediment samples were obtained in 21 stations, distributed along with 8 localities including different benthic habitats (Figure 1, Table 1), the bathymetric range of our samples fluctuated from 0 to 18 m. Samples were collected by digging surface sandy sediments in selected intertidal habitats and/or using PVC corers (5 cm internal diameter) through free diving. Local physical characteristics of benthic habitats, including the presence of biological components (e.g. macroalgae), type of sediment (biogenic, mixed, etc.) were registered. The samples were stored *in situ* in plastic jars and buckets. No fixative chemicals were used, live specimens were employed in all cases to avoid critical changes in key morphological traits used in the taxonomical identification.

**Table 1.**
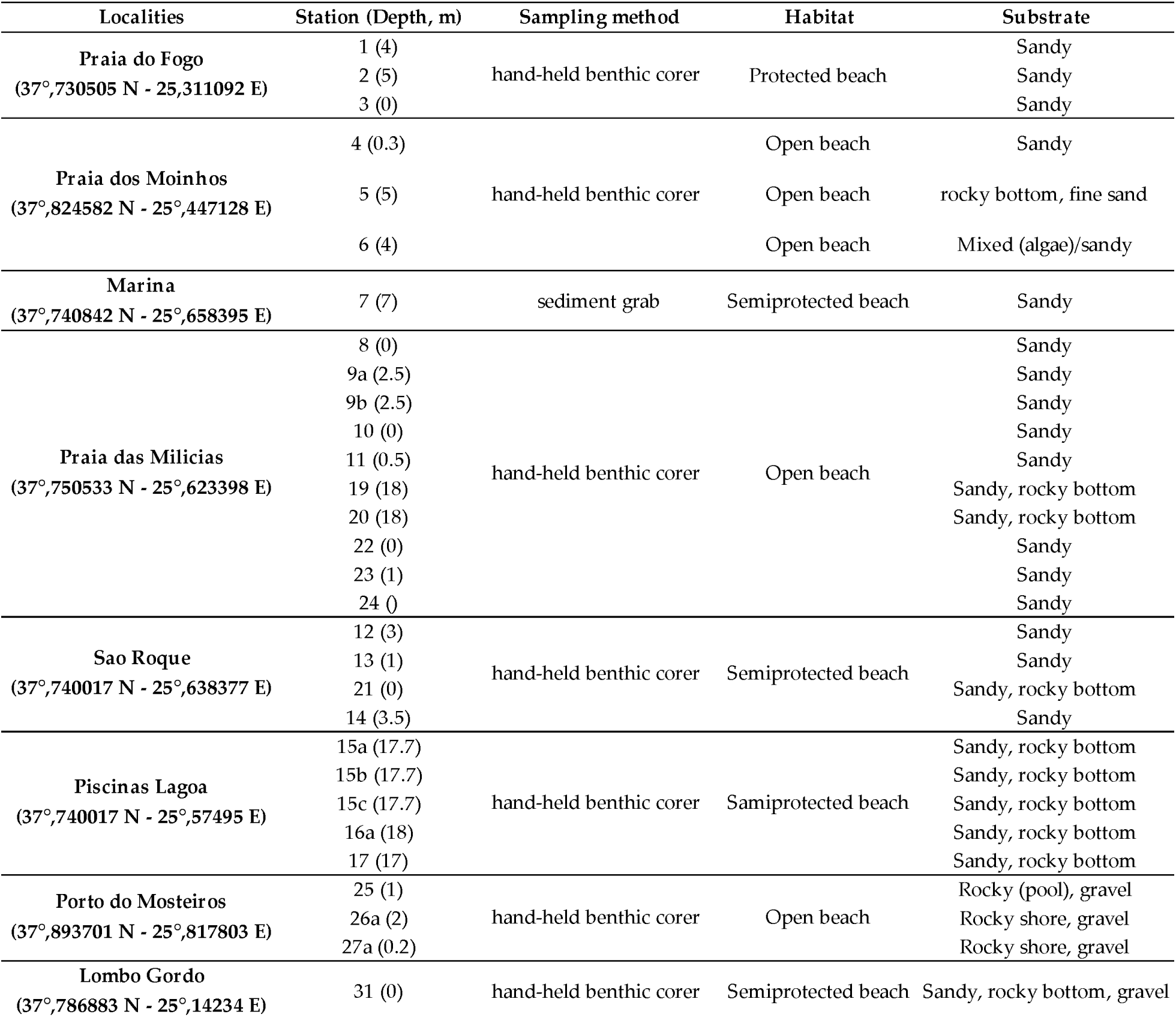
Main information of localities studied, habitats, sampling methods and the number of stations employed during the study.

**Figure 1.**
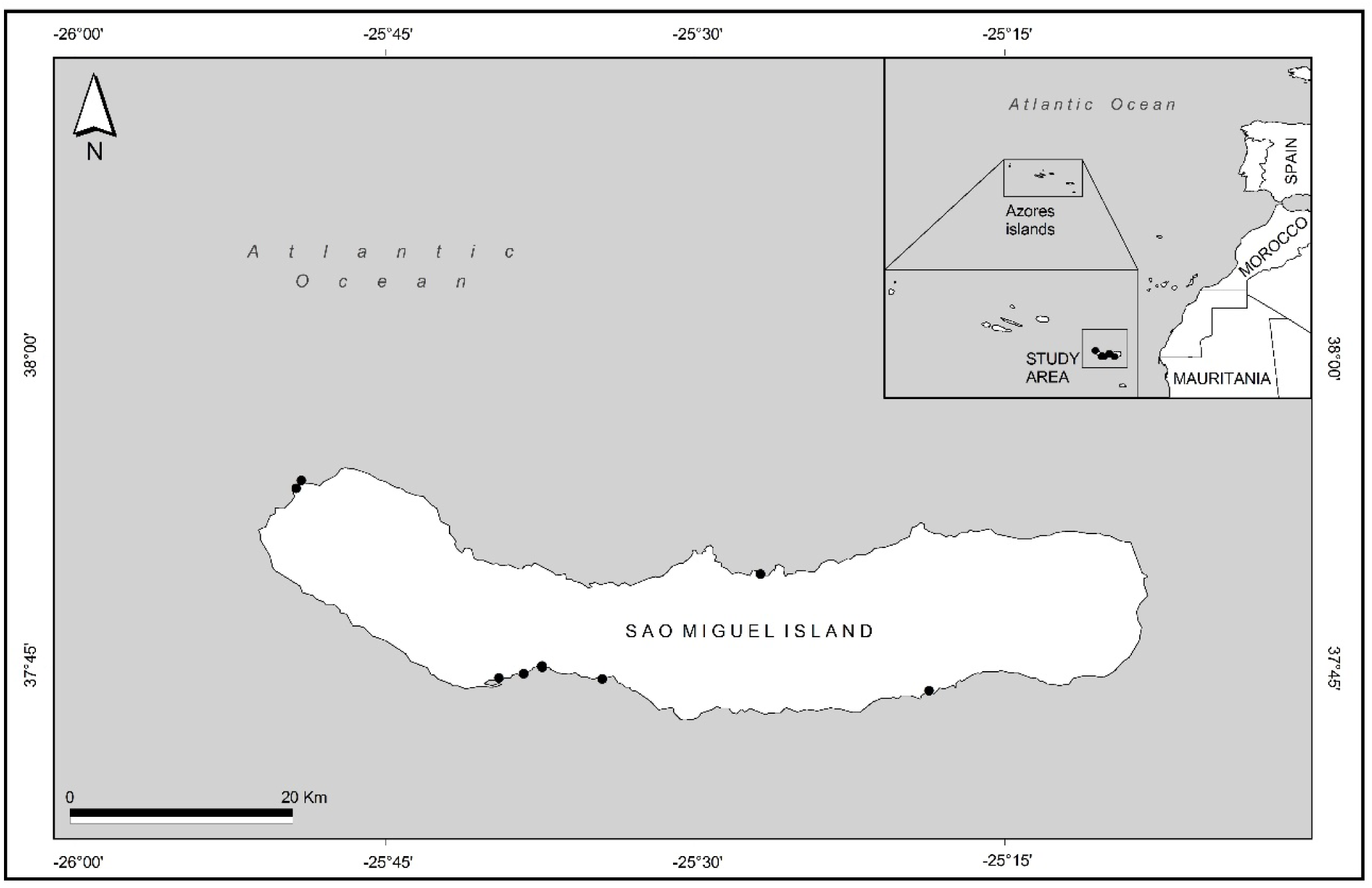
Location of sampling stations at São Miguel island, Azores archipelago, Portugal.

In the laboratory, meiofauna was extracted from the sediment by decantation, with some modifications in the procedure employed (see Pfannkuche & Thiel, 1988; Vincx, 1996). Meiofaunal organisms retained in a 35 µm sieve were then used for morphological characterization. Nematode specimens were manually separated from meiofauna bulk samples using a dissecting scope Nikon, mounted in glass slides with glycerin, and identified with a compound microscope (100X) Nikon Eclipse E1000, using pictorial keys (Platt & Warwick, 1983; 1988; Warwick *et al*., 1998), and the Nemys database (Becerra *et al*., 2019).

## Results

The nematode fauna at São Miguel island was represented by 4 orders, 21 families, and 43 genera. Enoplida had 8 families and 13 genera, while Chromadorida was represented by showed 7 families and 18 genera, thus being the most diverse nematode order. Monhysterida had five families and 10 genera, whereas Araeolaimida was only represented by 1 family and 2 genera (Fig. 2).

**Figure 2.**
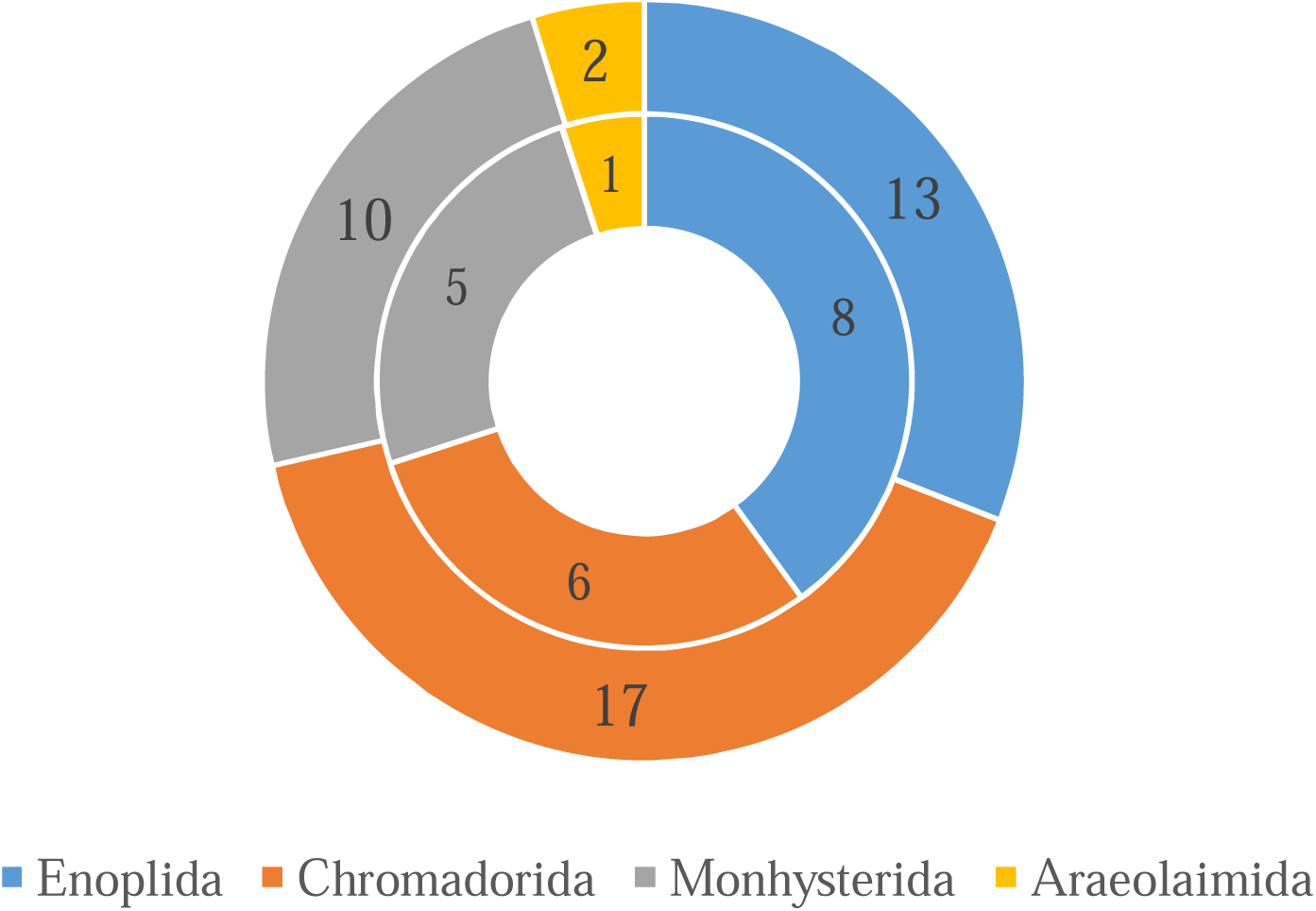
Taxonomic composition of free-living marine nematodes (Order rank) at São Miguel island, Azores archipelago, Portugal.

A total of 226 specimens were identified to the genus level (Fig. 3). The most abundant genera, which altogether account for 75% of the nematode fauna, included *<u>Adoncholaimus</u>* (32 individuals), *<u>Axonolaimus</u>* (18), *<u>Cyatholaimus</u>* (17), *<u>Enoploides</u>* (13), *<u>Rhabdocoma</u>* and *<u>Acanthopharynx</u>* (both with 11). The genera *<u>Viscosia</u>* and *<u>Enoplolaimus</u>* were represented by 7 specimens, while *<u>Halalaimus</u>, <u>Desmoscolex</u>, <u>Monophostia</u>, <u>Daptonema</u>*, and *<u>Theristus</u>* only by 6 specimens (Fig. 4). Most likely, some of the specimens identified in this study represent undescribed nematode. *<u>Cyatholaimus</u>, <u>Desmodora</u>* and *<u>Daptonema</u>* had two morphospecies each one.

**Figure 3.**
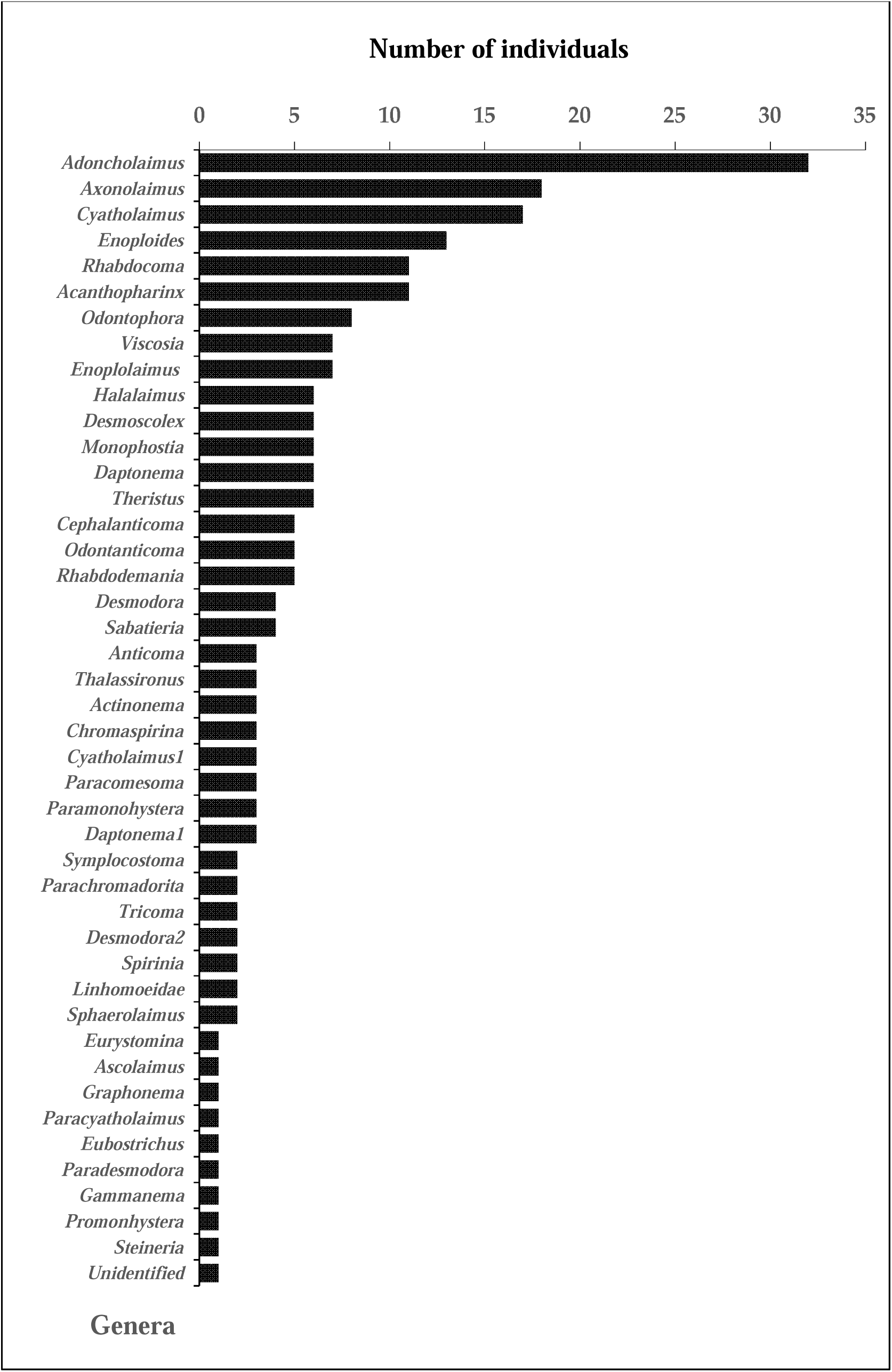
Abundance of free-living marine nematode taxa at São Miguel island, Azores archipelago, Portugal.

**Figure 4.**
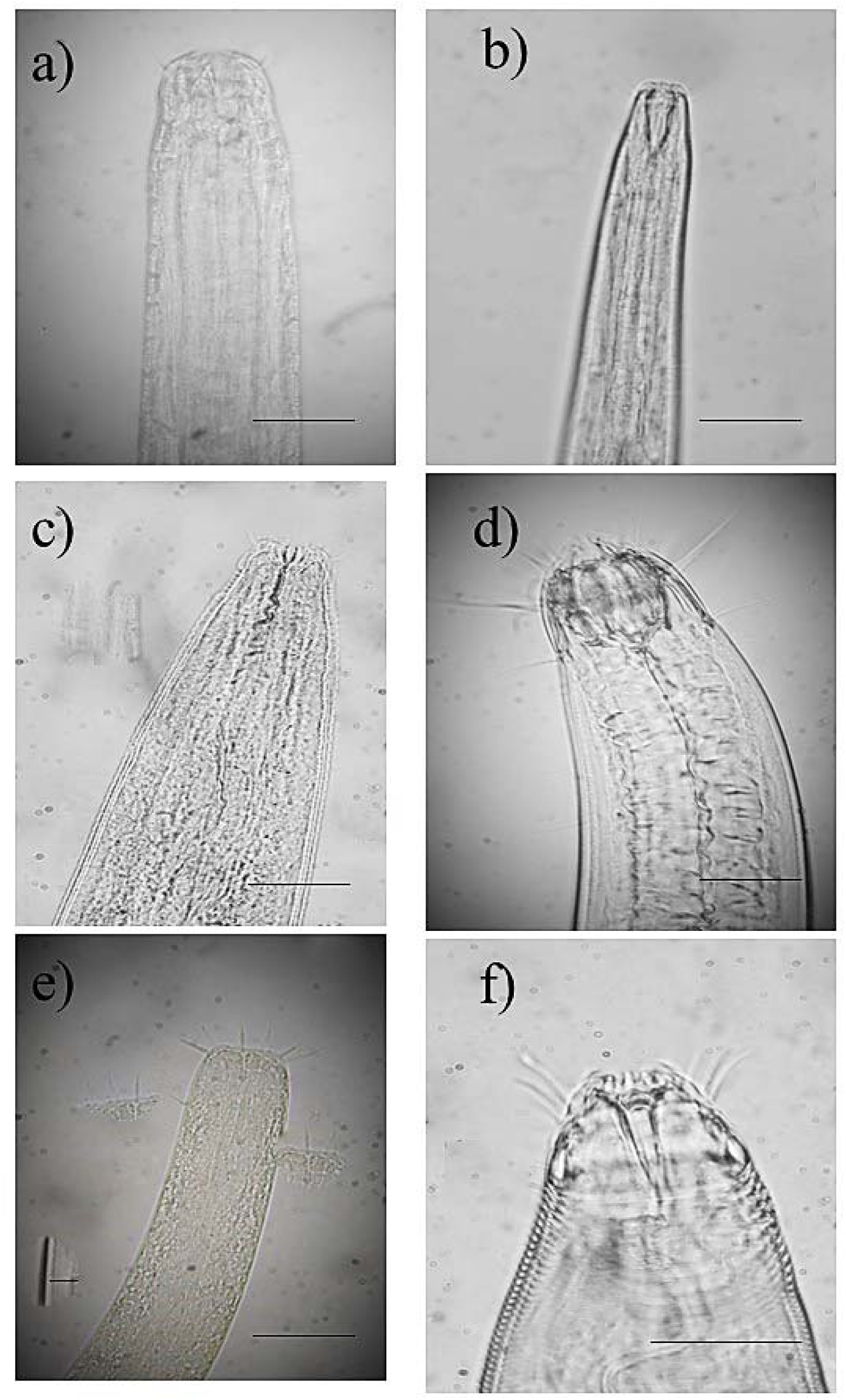
Anterior region (i.e. mouthparts) of selected marine nematode specimens. These genera were considered the six most abundant nematodes in the shallow benthic habitats studied at São Miguel island, Azores archipelago, Portugal. a) *Adoncholaimus*, b) *Axonolaimus*, c) *Cyatholaimus*, d) *Enoploides*, e) *Rhabdocoma*, and f) *Acanthoparynx*. Scale bars: 50 µm.

Sites with the highest nematode abundances included station 16 (40 specimens) and station 19 (36 specimens), followed by stations 15a and 15b, with 26 and 22 specimens, respectively. Station 7 had 13 organisms, while station 4 had 10 specimens. The rest of the stations revealed less than 10 organisms (Table 2).

**Table 2.** Preliminary list of free-living marine nematodes from shallow benthic habitats at Isla São Miguel, Azores.

| Taxon | Stations |  |  |  |  |  |  |  |  |  |  |  |  |  |  |  |  |  |  |  |  | Total |
| --- | --- | --- | --- | --- | --- | --- | --- | --- | --- | --- | --- | --- | --- | --- | --- | --- | --- | --- | --- | --- | --- | --- |
|  | 3 | 4 | 5 | 6 | 7 | 8 | 9a | 11 | 12 | 13 | 15a | 15b | 15c | 16 | 17 | 19 | 20 | 21 | 23 | 26a | 27a |  |
| *Anticoma* |  |  |  |  |  |  |  |  | 3 |  |  |  |  |  |  |  |  |  |  |  |  | 3 |
| *Cephalanticoma* |  |  |  |  |  |  |  |  |  | 1 | 2 |  |  | 2 |  |  |  |  |  |  |  | 5 |
| *Odontanticoma* |  |  |  |  |  |  |  |  |  |  |  |  |  |  |  | 5 |  |  |  |  |  | 5 |
| *Eurystomina* |  |  |  |  |  |  |  |  |  |  |  |  |  | 1 |  |  |  |  |  |  |  | 1 |
| *Symplocostoma* |  |  |  |  |  |  |  |  |  |  | 1 |  |  | 1 |  |  |  |  |  |  |  | 2 |
| *Thalassironus* |  |  |  |  |  |  |  |  |  |  | 1 |  |  |  |  | 1 | 1 |  |  |  |  | 3 |
| *Adoncholaimus* |  |  | 6 | 4 |  |  |  |  | 1 | 3 | 7 |  | 3 | 8 |  |  |  |  |  |  |  | 32 |
| *Viscosia* |  |  |  |  |  |  |  | 2 |  |  | 3 |  |  | 1 |  |  | 1 |  |  |  |  | 7 |
| *Halalaimus* |  |  |  |  |  |  |  |  |  |  |  | 6 |  |  |  |  |  |  |  |  |  | 6 |
| *Rhabdodemanina* |  |  |  |  |  |  |  |  |  |  |  | 2 | 1 | 1 |  | 1 |  |  |  |  |  | 5 |
| *Enoploides* |  |  |  |  |  |  |  |  |  |  | 1 |  |  | 5 |  | 2 |  |  | 4 |  | 1 | 13 |
| *Enoplolaimus* |  |  |  | 4 |  |  |  |  | 1 |  |  |  |  |  |  |  |  |  | 2 |  |  | 7 |
| *Rhabdocoma* |  |  |  |  |  |  |  |  | 1 |  | 1 | 5 |  | 2 |  |  |  |  | 2 |  |  | 11 |
| *Ascolaimus* | 1 |  |  |  |  |  |  |  |  |  |  |  |  |  |  |  |  |  |  |  |  | 1 |
| *Axonolaimus* |  |  |  |  |  |  |  |  |  |  | 4 | 1 |  | 6 |  | 8 |  |  |  |  |  | 18 |
| *Odontophora* |  |  |  |  |  |  |  |  |  |  |  |  |  | 3 |  | 5 |  |  |  |  |  | 8 |
| *Actinonema* |  |  |  |  |  |  |  |  |  |  |  | 2 |  |  |  |  |  |  |  |  | 1 | 3 |
| *Chromaspirinia* |  | 3 |  |  |  |  |  |  |  |  |  |  |  |  |  |  |  |  |  |  |  | 3 |
| *Graphonema* |  |  |  |  |  |  |  |  |  |  | 1 |  |  |  |  |  |  |  |  |  |  | 1 |
| *Parachromadorita* |  | 2 |  |  |  |  |  |  |  |  |  |  |  |  |  |  |  |  |  |  |  | 2 |
| *Cyatholaimus sp1* |  | 5 |  |  |  | 3 |  |  |  |  |  |  |  | 2 |  |  |  | 5 |  |  | 2 | 17 |
| *Cyatholaimus sp2* |  |  |  |  |  |  |  |  | 1 |  |  |  |  | 1 |  |  |  | 1 |  |  |  | 3 |
| *Paracyatholaimus* |  |  |  |  |  |  |  |  |  |  |  |  |  | 1 |  |  |  |  |  |  |  | 1 |
| *Desmoscolex* |  |  |  |  |  |  |  |  |  |  | 1 |  |  |  |  |  |  |  |  | 5 |  | 6 |
| *Tricoma* |  |  |  |  |  |  |  |  |  |  |  | 2 |  |  |  |  |  |  |  |  |  | 2 |
| *Acanthopharinx* |  |  |  |  |  |  |  |  |  |  | 3 | 2 | 1 | 1 |  | 3 | 1 |  |  |  |  | 11 |
| *Desmodora sp1* |  |  | 2 |  |  |  |  |  |  |  |  |  |  |  |  | 2 |  |  |  |  |  | 4 |
| *Desmodora sp2* |  |  |  |  |  |  |  |  |  |  |  |  |  | 1 |  |  |  |  | 1 |  |  | 2 |
| *Eubostrichus* |  |  |  |  |  |  |  |  |  |  |  |  |  |  |  | 1 |  |  |  |  |  | 1 |
| *Paradesmodora* |  |  |  |  |  |  |  |  |  |  |  |  |  | 1 |  |  |  |  |  |  |  | 1 |
| *Spirinia* |  |  |  |  |  |  |  |  |  |  |  |  |  |  |  | 2 |  |  |  |  |  | 2 |
| *Monophostia* |  |  |  |  |  |  |  |  | 1 |  | 1 |  |  | 2 |  |  |  | 2 |  |  |  | 6 |
| *Gammanema* |  |  |  |  |  |  |  |  |  |  |  |  |  | 1 |  |  |  |  |  |  |  | 1 |
| *Paracomesoma* |  |  |  |  | 3 |  |  |  |  |  |  |  |  |  |  |  |  |  |  |  |  | 3 |
| *Sabatieria* |  |  |  |  | 4 |  |  |  |  |  |  |  |  |  |  |  |  |  |  |  |  | 4 |
| *Paramonohystera* |  |  |  |  |  |  |  |  |  |  |  |  |  |  | 3 |  |  |  |  |  |  | 3 |
| *Promonohystera* |  |  |  |  |  |  |  |  |  |  | 1 |  |  |  |  |  |  |  |  |  |  | 1 |
| *Linhomoeidae* |  |  |  |  | 2 |  |  |  |  |  |  |  |  |  |  |  |  |  |  |  |  | 2 |
| *Sphaerolaimus* |  |  |  |  | 2 |  |  |  |  |  |  |  |  |  |  |  |  |  |  |  |  | 2 |
| *Daptonema sp1* |  |  |  |  | 2 |  | 2 |  |  |  |  | 1 |  |  |  |  | 1 |  |  |  |  | 6 |
| *Daptonema sp2* |  |  |  |  |  |  |  |  |  |  | 1 | 1 | 1 |  |  |  |  |  |  |  |  | 3 |
| *Steineria* |  |  | 1 |  |  |  |  |  |  |  |  |  |  |  |  |  |  |  |  |  |  | 1 |
| *Theristus* |  |  |  |  |  |  |  |  |  |  |  |  |  |  |  | 6 |  |  |  |  |  | 6 |
| Unidentified |  |  |  |  |  |  |  | 1 |  |  |  |  |  |  |  |  |  |  |  |  |  | 1 |
| **Total** | **1** | **10** | **9** | **8** | **13** | **3** | **2** | **3** | **8** | **4** | **26** | **22** | **6** | **40** | **3** | **36** | **4** | **8** | **9** | **5** | **4** | **226** |

## Discussion

Shallow benthic habitats are part of marine ecosystems of high socioeconomic importance, and frequently vulnerable to human impacts. From an ecological aspect, people might see these areas (e.g. sandy beaches) as poorly diverse habitats, which somehow attracted very few ecological researches (Defeo and & McLachlan 2005). This is unfortunate since many shallow marine areas harbour numerous species, especially those belonging to the benthic meiofauna, which are important links between micro-fauna and macrofauna, participating actively in benthic food webs (Weslawski et al.*et al*., 2000).

This study reports for the first time a list of free-living marine nematodes living on shallow soft-bottom habitats for São Miguel island, hence for the Azores. Although our sampling effort was relatively low and therefore does not represent the entire nematode diversity of São Miguel island, we believe it is the first step in the right direction to understand the diversity of marine nematodes on the island and also gain insights of nematodes associated with remote volcanic islands. Despite we cannot directly compare the number of nematode morphospecies (43 genera and 21 families) recorded in São Miguel island with other sandy beaches around the world due to differences in sampling effort and beach morpho dynamic conditions, we observed that the nematode diversity of São Miguel island is similar (i.e., in the range) in terms of nematode composition, particularly some common Families, where between 30 and 179 species have been reported (Ott, 1972; Platt, 1977; Blome 1983; Nicholas, 2006; Maria *et al*., 2013; Semprucci *et al*., 2013; Bhadury *et al*., 2015). In the Canary island, Riera *et al*., (2014) reported 48 nematode species, with a dominance of *<u>Daptonema</u> <u>hirsutum</u>*, and *<u>Pomponema</u> <u>sedecima</u>*, other species present were *<u>Acanthopharynx</u> <u>aff</u> <u>denticulate</u>, <u>Actarjania</u>*, sp1 and *<u>Ceramonema</u> <u>yunfengi</u>* and *<u>Scaptrella</u> <u>cf</u> <u>cincta</u>*.

Our results are also similar to those reported by Maria *et al*. (2013) who found 54 genera belonging to 25 families in two sandy beaches of Brazil, higher than those reported by Bhadury *et al*. (2015) on the coasts of India (20 genera and 13 families), but lower than the diversity in Italy, where 55 genera and 21 families were reported, having Chromadoridae and Xyalidae as the most diverse families, with 9 and 7 genera, respectively (Semprucci *et al*., 2013), and lower than the Gulf of California where 80 genera were found (Mundo-Ocampo *et al*., 2007). Likewise, in a temperate sandy beach at the Belgian coast Gherskiere *et al*. (2004) found 65 nematode genera representing 26 families, where Xyalidae was the most diverse in the number of genera (11) and species (20).

For the sandy beaches of Italy, Semprucci *et al*. (2013) indicated that the most abundant families were Xyalidae (47%), Chromadoridae (14%), Axonolaimidae (9%), and Comesomatidae (8%). In our study, the dominant families were Oncholaimidae, Axonolaimidae, Cyatholaimidae, Thoracostomopsidae, and Desmodoridae, and closely resembles the findings of Maria *et al*. (2013) who found Chromadoridae, Cyatholaimidae, Desmodoridae, and Oncholaimidae as the dominant nematode families on sandy beaches in Brazil. In addition to these same nematode families, Bhadury *et al*. (20015) reported the presence of Oxystominidae, Chromadoridae and Sphaerolaimidae in India.

Even though our results are still preliminary, we can infer that the presence of some genera indicates environmental conditions. For example, in station 7 (the “Marina”) we found a typical nematode community of fine sediments, with a dominance of *<u>Paracomesoma</u>, <u>Sabatieria</u>* (both Comesomatidae), and *<u>Daptonema</u>* (Xyalidae). These nematode genera are considered indicators of high concentrations of organic matter (Semprucci *et al*., 2015; Bhadury *et al*., 2015). As the first checklist of marine nematodes for São Miguel Island, our study showed that nematode diversity (i.e., regarding the number of genera and families) is within the range found in other sandy beach habitats. Furthermore, our study found congruence concerning the dominant families often found in sandy beaches. Additional information related to the type and origin of sediments is needed to fully understand the distribution patterns of nematode communities in São Miguel island. In fact, sediments in this remote island are predominantly of volcanic origin and are certainly an important environmental factor structuring nematode communities in this region. In Trindade, Brazil, Santos & Venekey (2017) found that nematode fauna was composed mainly of non-selective deposit feeders, with a total of 27 genera from 12 families, and Cyatholaimidae, Xyalidae and Oncholaimidae as the most abundant, the same families present in São Miguel. Therefore, we must consider it when making comparisons between nematode communities of sandy beaches. Nevertheless, we believe that this preliminary dataset on marine nematode communities will serve as a baseline for future taxonomic and ecological research in São Miguel as well as other volcanic islands.

Our results are also interesting because soft-sediment infauna (< 500 μm) in the Azores has been regarded as impoverished, particularly in shallow water sites (< 20 m in depth; e.g., Morton *et al*., 1998; Bamber & Robbins, 2009). This low infauna diversity at the Azores has been related to sediment instability (Ávila *et al*., 2008; Bamber & Robbins, 2009). Benthic soft-bottom habitats in the Azores are particularly devoid of macroinvertebrates, thus highlighting the importance of studying other infaunal taxa such as marine nematodes for environmental monitoring programs. Moreover, nematode communities could also be used as an indicator for Ecological Quality Status assessments in the scope of the water framework directive, similarly to what has been advocated for the Mediterranean (e.g., Moreno *et al*., 2011; Semprucci *et al*., 2015). Before it becomes a reality, however, additional work is needed to be put forward for a more comprehensive study (e.g., incorporating temporal variation) of marine nematode communities in the Azores archipelago.

### Concluding remarks

Our study shows the first results regarding the local diversity of free-living nematodes and provides scientific evidence about the diversity of small invertebrates (generally poorly studied or unconsidered in environmental studies). It also poses the need for increased efforts to improve the taxonomic analysis of particularly adaptive and diverse groups such as meiofauna. Moreover, as observed in other similar study areas, there is a significant possibility to suspect that many local benthic species (including some species here reported) could be new for science and might be endemic and/or restricted to local oceanographic conditions. On the other hand, our results also show that even under a limited sampling effort, these benthic communities can exhibit high diversity, possibly suggesting an ability to successfully occupy different types of shallow benthic habitats and use of available resources. More efforts are needed to understand the ecological role and value of these species on highly dynamic, shallow environments, and their changes over time.

## Acknowledgements

AJN, VA, and AMD thank Dr Katharina Jörger, the VW Foundation, and DRCT M3.3.B/ORG.R.C./020/2019 (funding) for facilitating our participation in the Summer Meiofauna Workshop. Thanks to the meiofauna team for sorting nematode specimen and to the University of the Azores for allowing us to use laboratories and the logistical help provided.

## Authors’ contribution statement

AJN and VA identified nematode specimens and wrote the manuscript; AMD processed sediment samples; ACC helped with laboratory equipment and wrote the manuscript.

